# LAT1 enables T cell activation under inflammatory conditions: a new therapeutic target for rheumatoid arthritis

**DOI:** 10.1101/2022.12.20.520910

**Authors:** Joy Ogbechi, Helen L. Wright, Stefan Balint, Louise M. Topping, Zec Kristina, Yi-Shu Huang, Eirini Pantazi, Maarten Swart, Dylan Windell, Eros Marin, Michael F. Wempe, Hitoshi Endou, Andrew M. Thomas, Andrew Filer, Trevor W. Stone, Alexander J. Clarke, Michael L. Dustin, Richard O. Williams

## Abstract

**Objective:** To assess the L-type amino acid transporter-1 (LAT1) as a possible therapeutic target for rheumatoid arthritis (RA).

**Methods:** Synovial LAT1 expression was monitored by immunohistochemistry and transcriptomic datasets. The contribution of LAT1 to gene expression and immune synapse formation was assessed by RNA-sequencing and total internal reflection fluorescent (TIRF) microscopy, respectively. Mouse models of RA were used to assess the impact of therapeutic targeting of LAT1.

**Results:** LAT1 was strongly expressed by CD4^+^ T cells in the synovial membrane of patients with active RA and the level of expression correlated with levels of ESR and CRP as well as DAS-28 scores. Deletion of LAT1 in murine CD4^+^ T cells inhibited the development of experimental arthritis and prevented the differentiation of CD4^+^ T cells expressing IFN-γ and TNF-α, without affecting regulatory T cells. LAT1 deficient CD4^+^ T cells demonstrated reduced transcription of genes associated with TCR/CD28 signalling, including *Akt1, Akt2, Nfatc2, Nfkb1* and *Nfkb2*. Functional studies using TIRF microscopy revealed a significant impairment of immune synapse formation with reduced recruitment of CD3ζ and phospho-tyrosine signalling molecules in LAT1 deficient CD4^+^ T cells from the inflamed joints but not the draining lymph nodes of arthritic mice. Finally, it was shown that a small molecule LAT1 inhibitor, currently undergoing clinical trials in man, was highly effective in treating experimental arthritis in mice.

**Conclusions:** It was concluded that LAT1 plays a critical role in activation of pathogenic T cell subsets under inflammatory conditions and represents a promising new therapeutic target for RA.

**Key Messages:** *What is already known about this subject?:*

- LAT1 is an amino acid transporter that has previously been shown to play a role in T cell activation.

*What does this study add?:*

- LAT1 is expressed by synovial T cells in human rheumatoid arthritis and the level of expression correlates with disease severity.
- LAT1 expression by T cells is necessary for development of severe arthritis in animal models.
- LAT1 is required for immune synapse formation and activation of pathogenic CD4^+^ T cell subsets in the inflamed joint, but not the lymph nodes.
- A small molecular weight LAT1 inhibitor, currently in clinical trials for cancer, is highly effective in animal models of rheumatoid arthritis.

*How might this impact on clinical practice of future developments?:*

- The context-specific nature of LAT1 involvement in T cell activation positions it as an ideal therapeutic target to distinguish between pathogenic and protective T cell responses and this study provides the scientific rationale for clinical evaluation of LAT1 inhibitors in the treatment of rheumatoid arthritis.

## INTRODUCTION

Pharmacological modulation of T helper cell activity currently involves the use of drugs targeting immune receptors or their downstream intracellular signalling pathways. However, there is a growing appreciation of the importance of metabolism-associated environmental cues in fine-tuning T cell activity and this provides novel opportunities for therapeutic intervention. T cells are critically dependent on uptake of amino acids (1), the availability of which exerts a powerful influence on T cell differentiation (2–5). Uptake of amino acids is regulated by specialised transporters, such as L-type amino acid transporter 1 (LAT1), also known as SLC7A5. LAT1 mediates sodium-independent influx of a range of large neutral amino acids (Leu, Val, Ile, Phe, Trp, His, Met, Tyr) (6) and exists in the plasma membrane as a heterodimeric complex covalently bound to CD98 (SLC3A2) (7). However, of the two subunits, LAT1 is solely responsible for the transport of amino acids (8) and is expressed by cells with a high nutrient demand (9), such as cancer cells (6, 10) and activated T and NK cells (11, 12).

Previous work has established the role of LAT1 in the proliferation and differentiation of CD4^+^ T cells (11) and in this study we set out to evaluate its contribution to the pathogenesis of rheumatoid arthritis (RA), a disease associated with dysregulated T helper cell responses. We first document the presence of LAT1 in the joints of RA patients and then relate its level of expression to clinical and serological markers of disease activity. We next set out to determine the role played by LAT1 in immune synapse formation and T cell activation at the site of inflammation in mice with experimental arthritis. Finally, we use animal models of arthritis to evaluate the efficacy of a small molecule LAT1 inhibitor currently undergoing clinical trials in man. Our findings reveal that LAT1 plays a non-redundant role in activation of specific CD4^+^ T cell subsets under inflammatory conditions and represents a highly tractable therapeutic target for RA.

## METHODS

See online supplemental material.

## RESULTS

### 1. LAT1 expression correlates with disease activity

Synovial tissue samples were collected during joint replacement surgery from patients with osteoarthritis (OA) or RA (diagnosed according to the 1987 American College of Rheumatology criteria (13)). LAT1 expression was detected by immunohistochemistry in 100% (7/7) of RA samples (Fig. 1A, B) in the lining, sublining and perivascular synovial regions, particularly at sites of lymphocytic infiltration (Fig. 1A). In contrast, LAT1 was detected in only 37% (3/8) of osteoarthritis (OA) samples, with a level of expression that was drastically reduced compared to RA (Fig. 1A). In RA synovium, LAT1 was detected mainly in CD4^+^ T cells and to a lesser extent in CD68^+^ myeloid cells (Fig. 1C).

**Fig 1:**
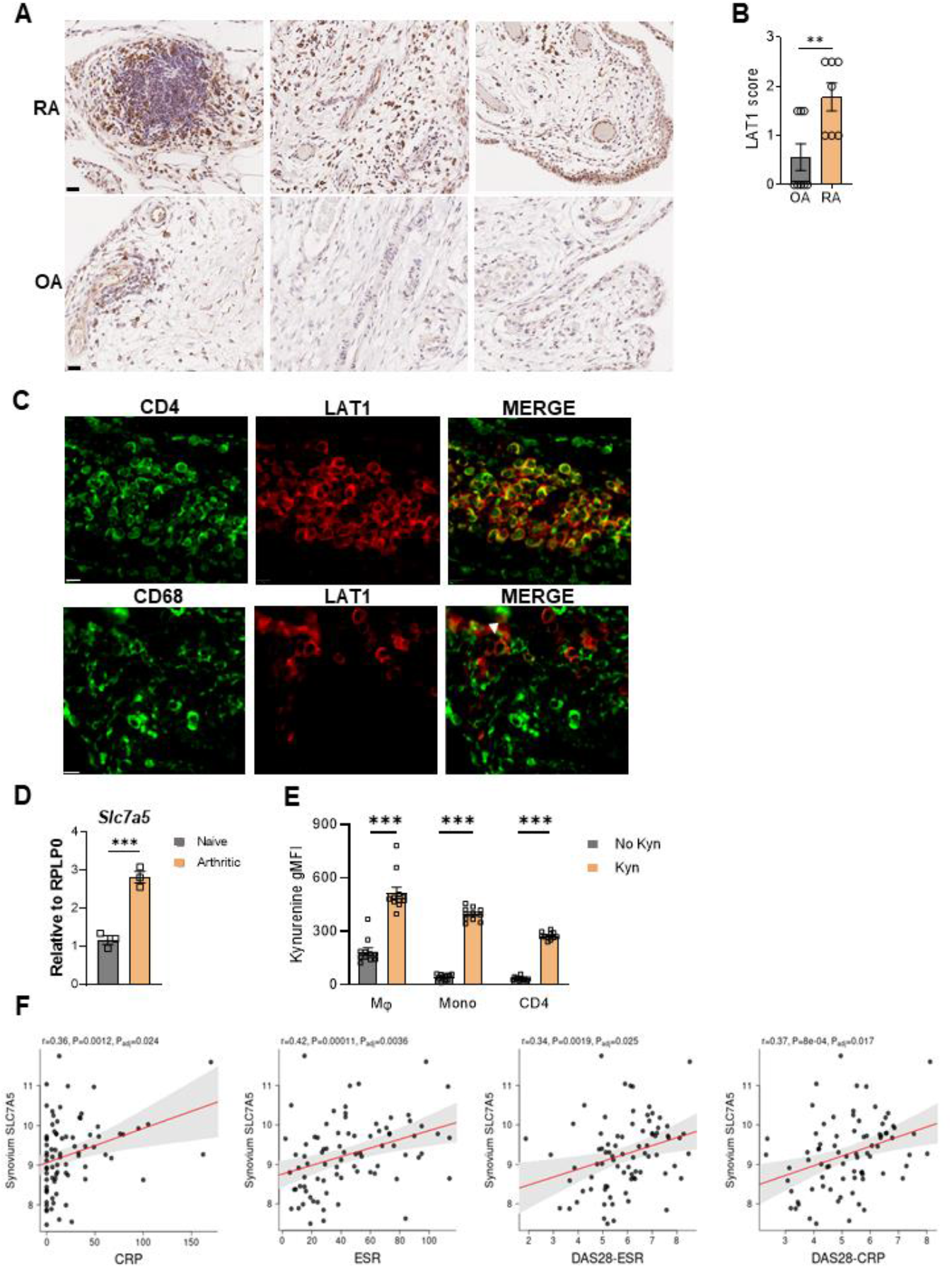
LAT1 expression correlates with disease activity. (A) Immunohistochemistry of human synovial tissue from RA (n=7) and OA (n=6) patients using anti-LAT1 antibody. Scale bar represents 20 μm (B) Immunohistochemistry staining score for LAT1 on human synovial tissue. (C) Double immunofluorescence staining of RA synovium using anti-LAT1 (red) and anti-CD4 or anti CD-68 (green) antibodies. Arrows indicate CD68^+^LAT1^+^cells, scale bar represents 10 μm. (D) qPCR analysis of *Slc7a5* mRNA levels in bulk knee samples from naïve mice and mice with AIA. (E) Kynurenine uptake assay performed on immune cells isolated from the knees of mice with AIA (mean±SEM, n=11, Mann-Whitney U test, from 3 independent experiments ***P < 0.001). (F) Two-tailed Spearman’s correlation between synovial expression of SLC7A5 with disease activity in the PEAC cohort (n = 90); grey-shaded area, 95% confidence interval; R and P values shown.

We then quantified *Slc7a5* RNA levels in knee joints of mice with antigen-induced arthritis (AIA). To induce arthritis, mice were immunised with methylated bovine serum albumin (mBSA), followed by intra-articular injection of mBSA. Compared with healthy joints, joints from arthritic mice showed a 3-fold increase in expression of *Slc7a5* (Fig. 1D). The functionality of LAT1 in CD4^+^ T cells, macrophages and monocytes from the joints of arthritic mice was confirmed by measuring uptake of kynurenine, a known LAT1 substrate that can be detected fluorometrically (14) (Fig. 1E, see Fig. S1B for gating strategy).

Lastly, to determine if LAT1 expression is associated with a particular disease pathotype and whether it correlates with disease activity, we analysed data from early, treatment-naïve, RA patients within the Pathobiology of Early Arthritis Cohort (PEAC) (15) (see methods). Expression of *SLC7A5* was significantly enriched in the lympho-myeloid RA pathotype which is rich in T cells, myeloid cells and CD20 B cell aggregates compared to the diffuse-myeloid pathotype (myeloid rich but poor in B cells; P_adj_=8×10^-6^) and fibroid pathotype (low immune-inflammatory cell infiltrate; P_adj_=4×10^-4^). Importantly, strong positive correlations were observed between synovial *SLC7A5* expression and the inflammatory markers: ESR (P_adj_=0.0036) and CRP (P_adj_=0.024), and the disease activity scores: DAS28-ESR (P_adj_=0.025) and DAS28-CRP (P_adj_=0.017) (Fig. 1G).

### 2. LAT1 expression in T cells is required for robust arthritis

The impact of LAT1 deletion in hematopoietic cells was assessed by crossing mice carrying floxed *Slc7a5* alleles (Slc7a5fl/fl) (11) with mice expressing Cre recombinase controlled by the *Vav* promoter (Fig 2A, B). Histological analysis of knee joints of mice with AIA revealed reduced cartilage damage and cell infiltration in LAT1^ΔVav^ mice compared with LAT1^WT^ mice (Fig 2C, D), indicating a significant role for LAT1 in development of arthritis.

**Fig 2:**
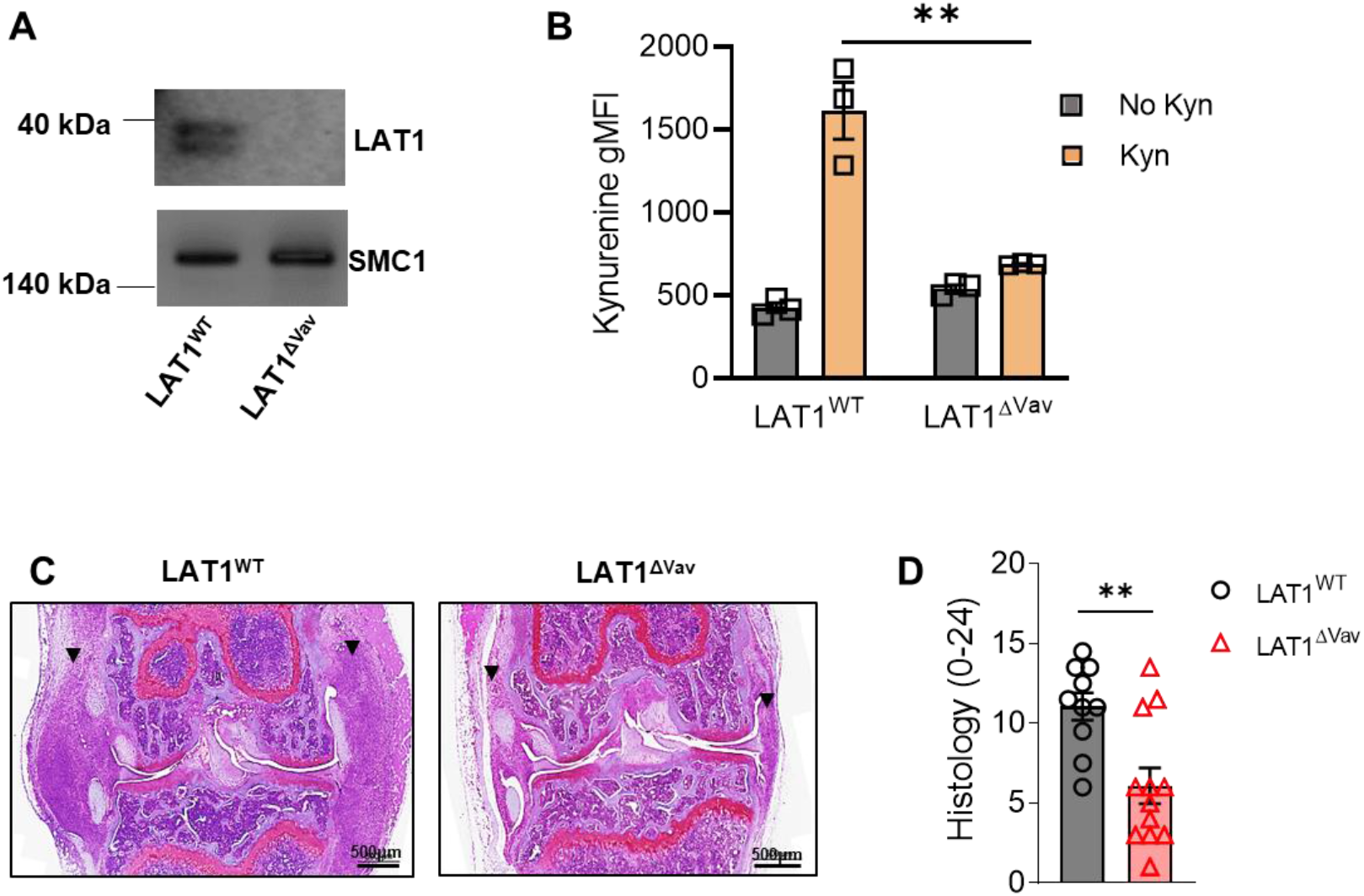
LAT1 deletion in hematopoietic cells inhibits development of AIA. (A) LAT1 expression detected by western blotting in CD4^+^ T cells isolated from LAT1^WT^ and LAT1^ΔVav^ mice 48h after TCR stimulation. (B) Kynurenine uptake assay performed on CD4^+^ T cells isolated from LAT1^WT^ and LAT1^ΔVav^ mice 48h after TCR stimulation. Bars indicate mean ± SD, n = 3 mice from one experiment representative of > 3 separate experiments. (C) Representative images of Safranin O stained knees obtained from mice 7 days post arthritis induction. (D) Histological analysis of arthritic knees (mean±SEM, n = 10-12, Mann-Whitney U test, from 3 independent experiments). (*P < 0.05; **P < 0.01, ***P < 0.001, 2-tailed unpaired Student t test.

We next sought to compare the impact of LAT1 deletion in myeloid *versus* lymphoid cells. Myeloid specific LAT1 deletion was achieved by crossing Slc7a5fl/fl mice with mice expressing Cre recombinase controlled by the LysM promoter (Fig S2). The development of arthritis was found to be unaffected by myeloid cell specific LAT1 deletion, with comparable disease severity and similar numbers of neutrophils, monocytes, macrophages and CD4^+^ T cells in LAT1^ΔLysM^ and LAT1^WT^ mice (Fig 3A–D). Similarly, no differences in numbers of cells expressing M1/M2 markers, MHCII, iNOS, CD206 and Arg-1 (Fig 3E) or cytokine levels in the joints were observed between LAT1^ΔLysM^ and LAT1^WT^ mice (Fig. 3F).

**Fig 3:**
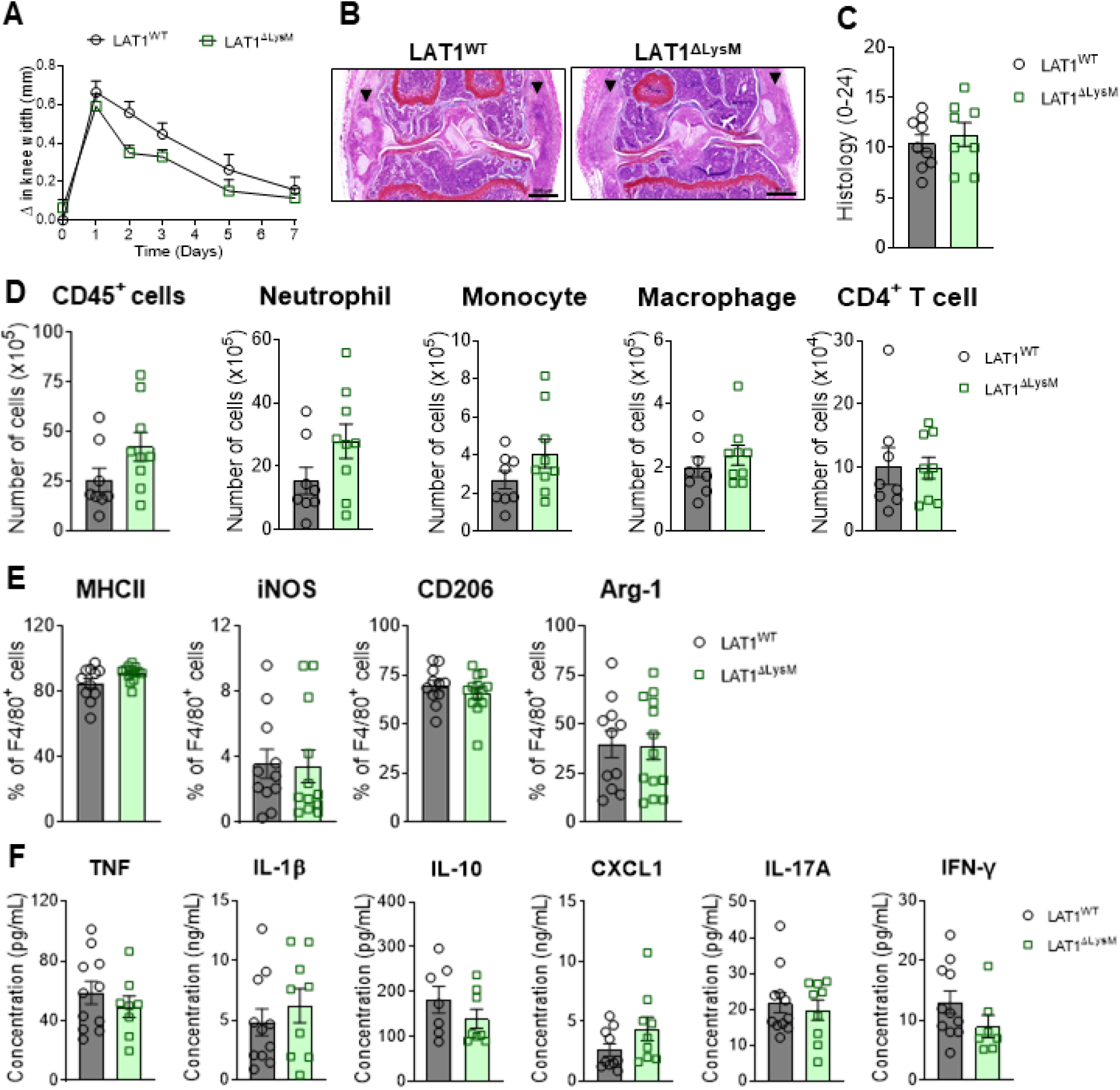
Deletion of LAT1 in myeloid cells does not affect arthritis. LAT1^WT^ and LAT1^ΔLysM^ mice were immunized with mBSA and challenged with mBSA or PBS, as a control, 21 days later. (A) Change in knee thickness following mBSA challenge (mean±SEM, n=9 from 3 independent experiments). (B) Representative images of Safranin O stained knees obtained from mice 7 days post arthritis induction. (C) Histological analysis of arthritic knees (mean±SEM, n=8-9 from 3 independent experiments). (D) Total number of immune cells (CD45^+^), neutrophils, monocytes, macrophages, and CD4^+^ T cells recovered from excised knees on day 5 post arthritis induction (mean±SEM, n=8-9 from 2 independent experiments). (E) Frequencies of macrophages (F4/80^+^CD64^+^) expressing the indicated proteins (mean±SEM, n=11-13 from 3 independent experiments). (F) Cytokine levels in the joint on day 5 post arthritis induction quantified by ELISA (mean±SEM, n=9-11 from two independent experiments).

To delete LAT1 in T cells, we crossed Slc7a5fl/fl mice with CD4-Cre mice (Fig S2A, B). Following arthritis induction, there was significantly less knee-swelling, cathepsin activity and pain in LAT1^ΔCD4^ mice compared to LAT1^WT^ mice (Fig 4A–D). Histological analysis confirmed reduced disease severity in LAT1^ΔCD4^ mice which was associated with reduced numbers of infiltrating immune cells and reduced expression of IL-1β, TNFα and IFNγ in the joints (Fig 4E–K). Further analysis of the joints and lymph nodes of arthritic LAT1^ΔCD4^ mice revealed reduced numbers of IFNγ^+^ CD4^+^ and TNF^+^CD4^+^ T cells compared to LAT1^WT^ mice, whilst numbers of CD25^+^ FoxP3^+^ CD4 T cells were unchanged (Fig 5A, 5B, 5D). Similarly, total numbers of IL-17^+^ CD4^+^ T cells in the lymph nodes and joints of LAT1^ΔCD4^ mice were unchanged (Fig 5C). There was no difference between LAT1^ΔCD4^ and LAT1^WT^ mice in the proportion of CD4^+^ T cells expressing the inhibitory receptors PD-1, CTLA4 and LAG-3 (Fig 5E) indicating that deletion of LAT1 does not lead to a state of T cell exhaustion, as has been described in chronic infection and cancer (16, 17).

**Fig 4:**
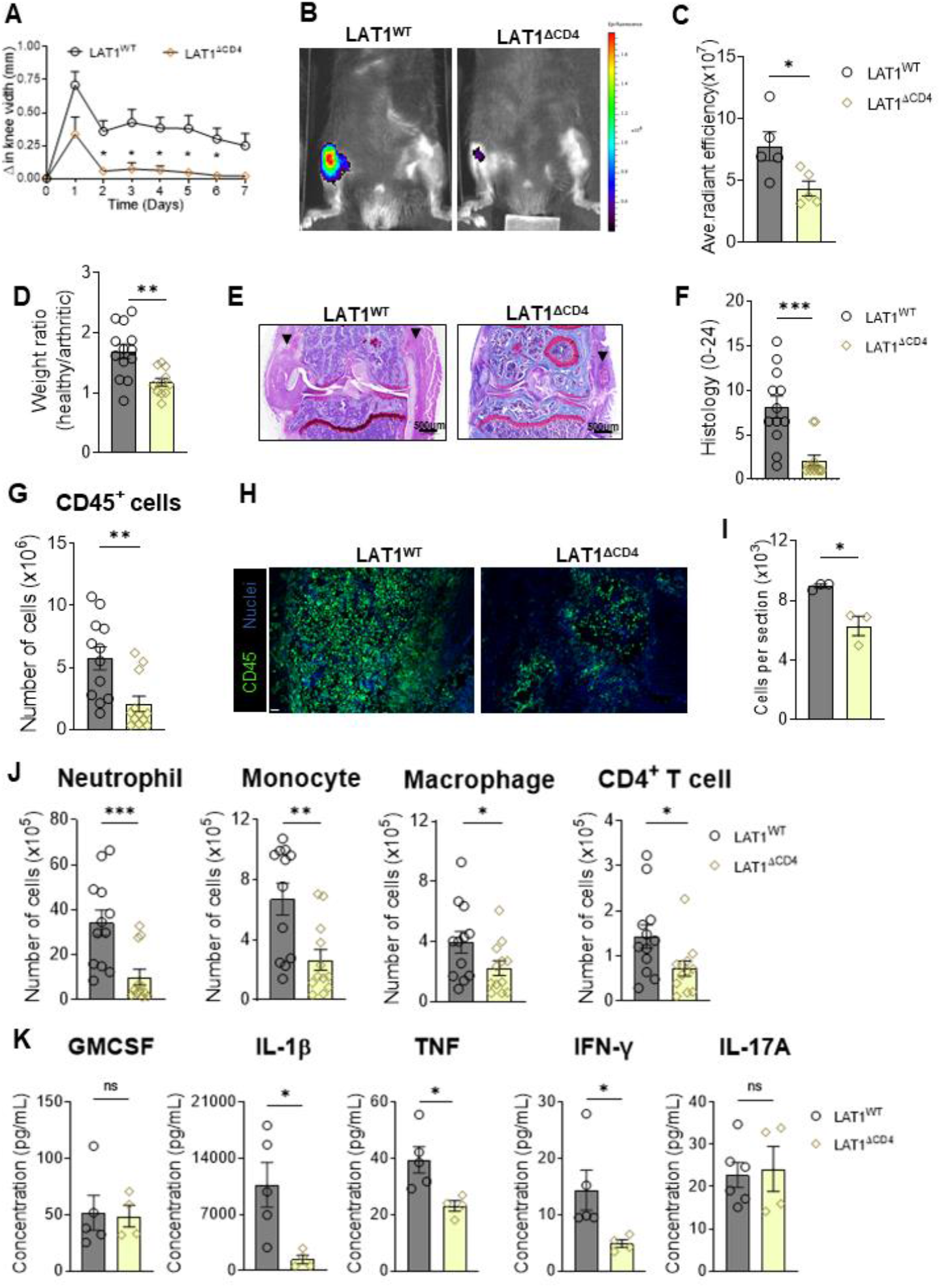
LAT1 expression in T cells is required for development of severe arthritis. AIA in LAT1^WT^ *versus* LAT1^ΔCD4^ mice. (A) Knee-swelling (mean±SEM; n=9; 2-way ANOVA with Tukey’s post hoc test). (B) Cathepsin activity (representative images). (C) Quantification of cathepsin activity (mean±SEM, n=5; Mann-Whitney U test). (D) Dynamic weight-bearing test (mean±SEM; n=11-13; Mann-Whitney U test). (E) Representative images of Safranin O stained knee joints. (F) Histological analysis (n=12; Mann-Whitney U test). (G) Total number of CD45^+^ cells recovered from knees on day 5 of arthritis (mean±SEM; n=12; Mann-Whitney U test). (H) Representative image of arthritic knee joint stained using SYTOX blue (nuclei, grey) and anti-CD45 (red). Scale bar represents 10 μm. (I) Quantification of CD45^+^ cells (mean±SD; n=3; Mann-Whitney U test). (J) Total number of neutrophils, monocytes, macrophages, and CD4^+^ T cells recovered from excised knees on day 5 of arthritis (mean±SEM; n=12; Mann-Whitney U test). (K) Cytokine levels in the knee on day 5 of arthritis (mean±SD; n=4-5; Mann-Whitney U test). *P<0.05; **P<0.01, ***P<0.001.

**Fig 5:**
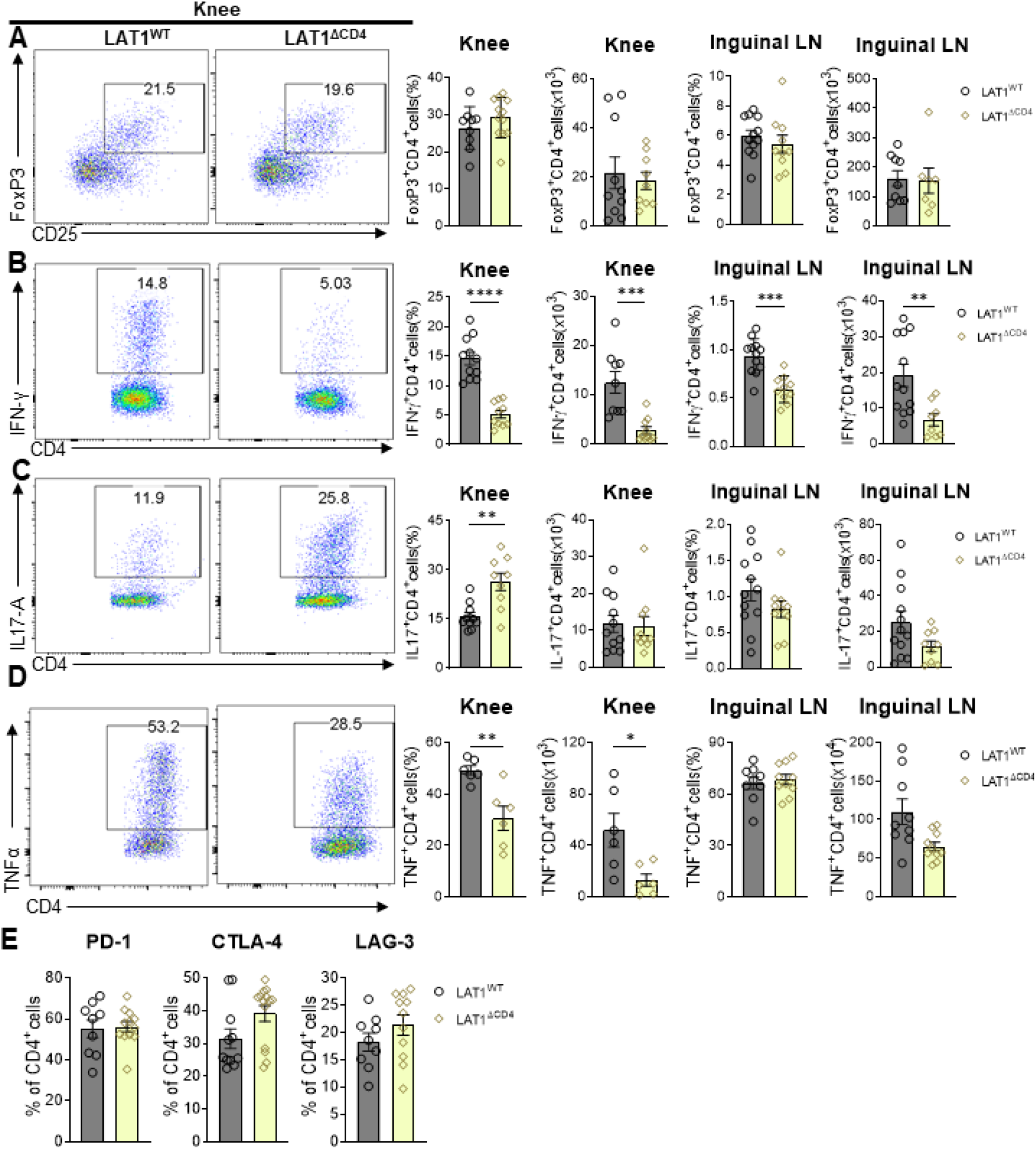
LAT1 deletion selectively impairs differentiation of IFNγ^+^ and TNF^+^ T cells. Cells were obtained from knee joints of LAT1^WT^ and LAT1^ΔCD4^ mice with AIA (day 5). Representative flow cytometric images of CD4^+^ T cells and frequencies and absolute numbers of (A) CD25 and Foxp3 double positive Tregs (B) IFN-γ^+^ Th1 cells (C) IL-17^+^ Th17 cells and (D) TNF^+^ T cells in the knee and draining lymph nodes. (E) Frequencies of PD-1, CTLA-4 and LAG-3 expressing CD4^+^ T cells in the knee (mean±SEM; n=6-12; Mann-Whitney U test). *P<0.05, **P<0.01, ***P<0.001.

### 3. Loss of LAT1 alters T cell transcriptional profile in arthritis

RNA-Seq analysis was carried out on CD4^+^ T cells from the arthritic joints of LAT1^WT^ and LAT1^ΔVav^ mice to identify pathways affected by LAT1 deletion. A large number of genes (4980 upregulated, 1207 downregulated) were altered in CD4^+^ T cells from LAT1^ΔVav^ mice (1.5-fold change in expression cut-off, Fig 6A). Consistent with our earlier findings, there were reductions in *Ifng* and *Tbx21* and *Tnf* in LAT1^ΔVav^ mice (Fig S3) whilst *Il2ra* and *Foxp3* were unaffected. Ingenuity Pathway Analysis (IPA) revealed down-regulation of multiple gene programs linked to cell cycle, T cell activation, TCR/CD28 signalling, T cell exhaustion, oxidative stress and apoptosis in LAT1 deficient CD4^+^ T cells (Fig 6A). IPA also predicted the down-regulation of a mechanistic network of genes regulated by activation of the T cell receptor complex, including CD28, CD3, IL-2, IFNɣ, and the transcription factors NF-κB, STAT1 and STAT3 (Fig 6B, P_adj_=7×10^-19^). Secondary validation by qPCR confirmed that LAT1 deletion results in decreased expression of *Akt1, Akt2, Nfatc2, Nfkb1, Nfkb2* (Fig 6C) which are linked to TCR/CD28 signalling, T cell exhaustion, apoptosis and NF-κB signalling pathways (Fig 6D).

**Fig 6:**
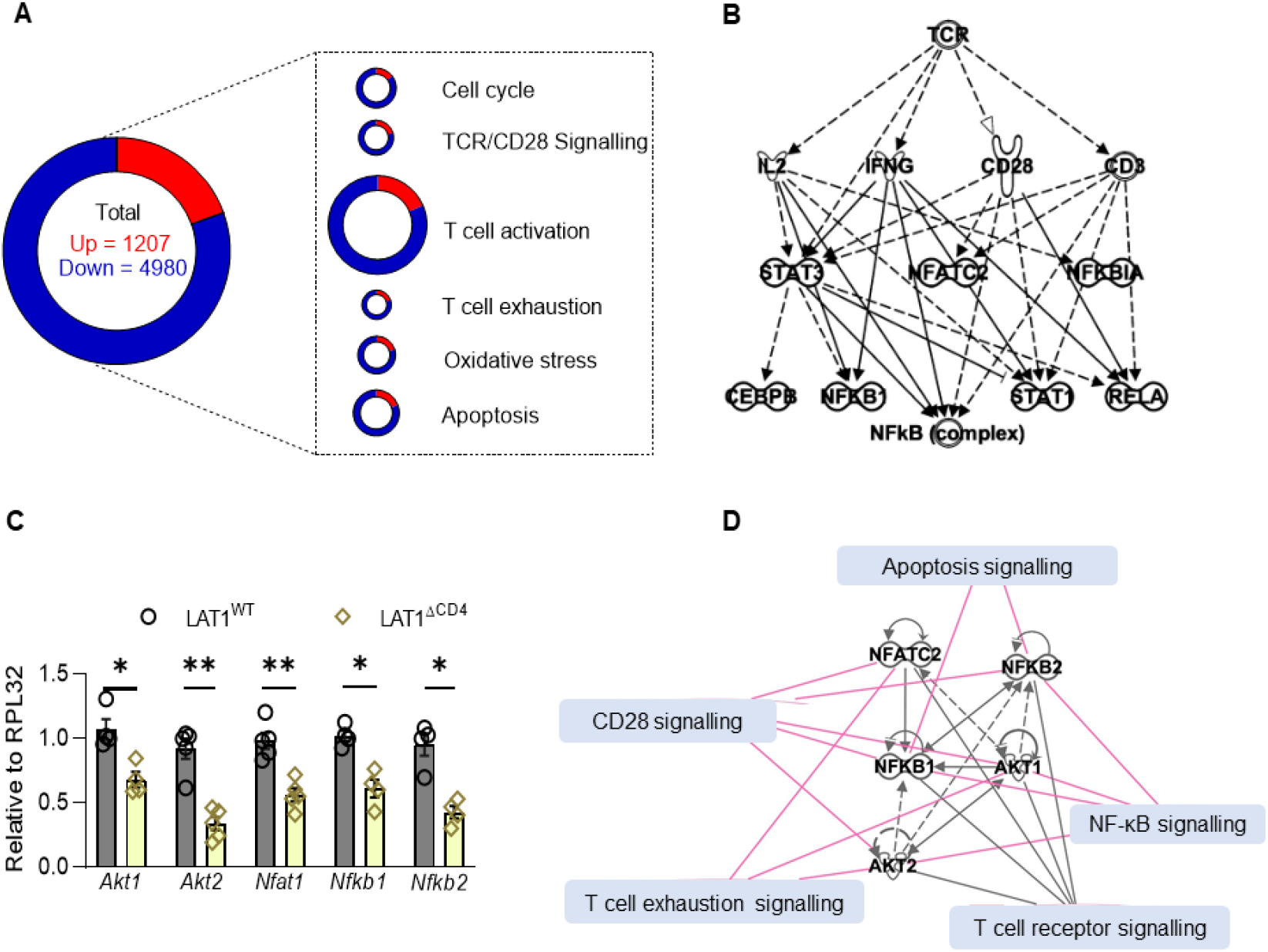
Transcriptome analysis of CD4^+^ T cells from knees of WT and LAT1^-/-^ mice with AIA. T cells were sorted from the knees (n=4) of arthritic LAT1^WT^ and LAT1^ΔVav^ mice and RNA was extracted for sequencing. (A) Categorisation of altered genes by IPA in LAT1 deficient mice. Circle size is proportional to the number of genes. (B) IPA predicted down-regulation of a mechanistic network of genes, receptors and transcription factors down-stream of the TCR. Arrows indicate direct (solid) or indirect (dashed) signalling inhibition. (C) Validation of genes regulated by LAT1 in CD4^+^ T cells by qPCR (mean±SD; n=4; unpaired t test, *P<0.05). (D) Cartoon showing interactions of genes shown in C and the canonical signalling pathways to which they are linked.

### 4. LAT1 regulates TCR signalling in the joint

We next compared the signaling behavior of LAT1-sufficient and LAT1-deficient T cells from the joints and lymph nodes of arthritic mice, including their ability to form immune synapses. CD4^+^ T cells were incubated on activating surfaces coated with anti-CD3, anti-CD28 and ICAM-1. Analysis of immune synapses of T cells from the joints by total internal reflection fluorescent microscopy (TIRFM) revealed reduced mean fluorescence intensity (MFI) of CD3ζ, a critical signaling molecule, and pTyr in LAT1^ΔCD4^ CD4^+^ T cells compared to LAT1^WT^ T cells (Fig 7A). Furthermore, the accumulation of the underlying actin, a cytoskeleton structural molecule required for immune synapse reorganization and stability which is also indicative of cell activation, was reduced in the CD4^+^ T cells from LAT1^ΔCD4^ versus LAT1^WT^ mice (Fig 7E) indicating that LAT1 deletion in arthritis reduces the intensity of TCR triggering. In contrast, quantification of the TIRFM images revealed equal levels of activation markers from CD4^+^T cells isolated from the lymph node of LAT1^WT^ and LAT1^ΔCD4^ mice (Fig 7F). Cells spread equally on the ICAM-1 surface (Fig 7G) and the CD3ζ and the pTyr levels were not significantly different between LAT1^WT^ and LAT1^ΔCD4^ mice (Fig 7H–I). Similarly, the underlying actin skeleton was not altered (Fig 7J). This suggests that the reduced TCR activation in CD4^+^T cells from LAT1^ΔCD4^ mice only occurs under conditions of active inflammation in the knee.

**Fig 7:**
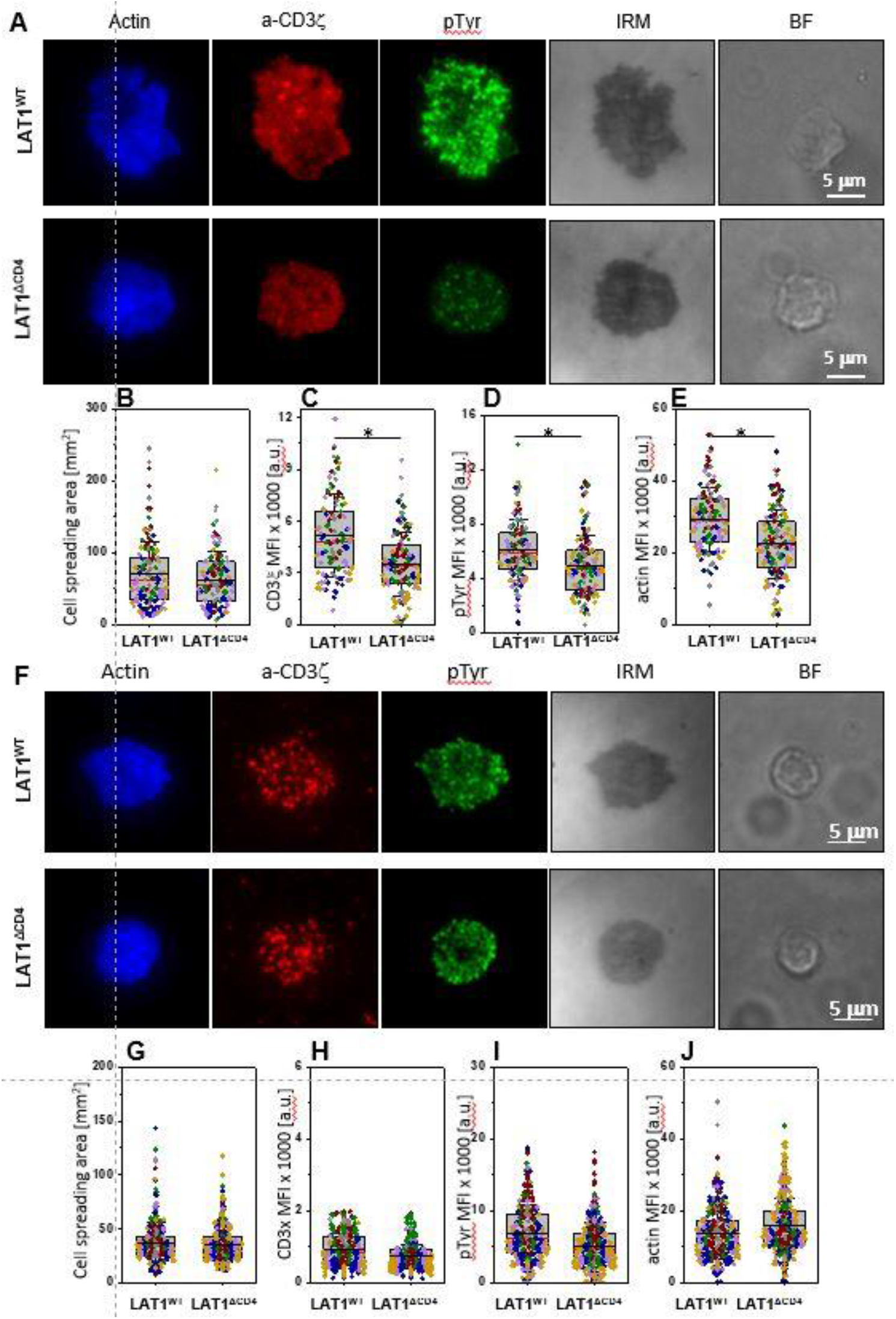
LAT1 regulates TCR signalling in joints but not lymph nodes, (A,F) Representative TIRFM images of CD4^+^ T cells from the joints (A) or lymph nodes (F) of arthritic LAT1^WT^ and LAT1^ΔCD4^ mice interacting with activating (ICAM1+anti-CD3+anti-CD28) surface for 15 mins. Cells were permeabilized and stained with antibodies against CD3ζ-AF647 (red), pTyr-AF488 (green). Actin cytoskeleton was labelled with phalloidin-CF568 (blue). IRM, interference reflection microscopy; BF, bright field. (B,G) Quantification of cell spreading area of T cells from joints (B) or lymph nodes (G) of LAT1^WT^ and LAT1^ΔCD4^ mice on activating surface. (C,H) Corresponding MFI of anti-CD3ζ. (D,I) anti-pTyr (D) and actin (E,J) as in A and F. Horizontal lines and error bars represent mean±SD (black line) and median (red line). Gray box represents 25-75 percentile. Data are from n=64-167 cells from 3-6 mice; each dot represents a cell; each colour represents a mouse. Two-tailed *t* test assuming unequal variances. *, P<0.05.

### 5. Deletion or inhibition of LAT1 ameliorates established arthritis

Finally, we assessed the therapeutic utility of systemic inhibition of LAT1 by pharmacologic inhibition or global genetic deletion. JPH203 (O-[(5-Amino-2-phenyl-7-benzoxazolyl)methyl]-3,5-dichloro-L-tyrosine dihydrochloride) is a selective LAT1 inhibitor currently undergoing clinical evaluation in humans (18, 19) and we first confirmed its ability to modulate T cell activity. Treatment of naïve CD4^+^ T cells with JPH203 *in vitro* (10 μM) inhibited the differentiation of Th1 and Th17 cells but not Treg cells (Fig S4A). Next, mice with established CIA or AIA were injected intraperitoneally with vehicle or JPH203 (50mg/kg/day). Clinical and histological analysis revealed that LAT1 inhibition caused a significant reduction in arthritis severity in both CIA (Fig 8A–C) and AIA (Fig D-G). Mice were monitored by a treatment blinded observer and no adverse effects were observed following administration of JPH203, such as weight loss, reduced movement or social isolation.

**Fig 8:**
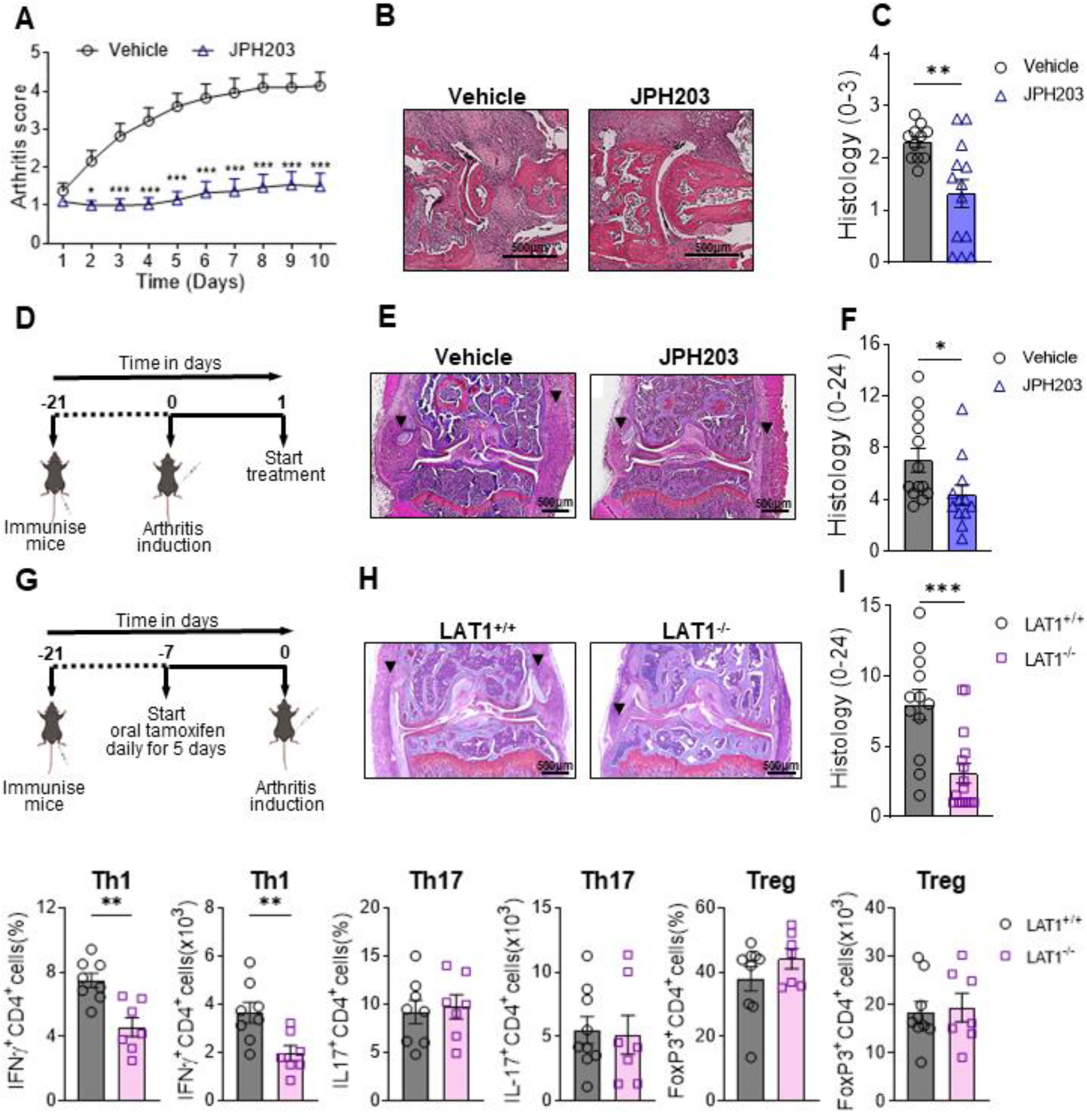
Inhibition of LAT1 attenuates established arthritis. (A-C) Mice with CIA were treated daily with JPH203 (50 mg/kg/day) or vehicle (A) Clinical scores (mean±SEM; n=24; 2-way ANOVA with Tukey’s post hoc test). (B) Representative images of proximal interphalangeal joint from mice treated with JPH203 or vehicle (C) Histological scores (mean±SEM; n=11-13; Mann-Whitney U test) (D) Treatment regimen for AIA. JPH203 was administered at 50 mg/kg/day. (E) Representative images of Safranin O stained knees joints. (F) Histological analysis (mean±SEM; n=12-13; Mann-Whitney U test). (G) Cartoon representation of regimen for tamoxifen administration to delete LAT1 in AIA model. (H) Representative images of Safranin O stained knees from mice on day 7 of arthritis. (I) Histological analysis of arthritic knees (mean±SEM; n=12-16; Mann-Whitney U test). (J) On day 5 of arthritis, cells were obtained from the knee and analysed by flow cytometry to determine frequencies and absolute numbers of T cell subsets (mean±SEM; n=7-9; Mann-Whitney U test). *P<0.05, **P<0.01, ***P<0.001.

We also assessed the effect on AIA of systemic deletion of *Slc7a5* in adult UbCre^+/-^Slc7a5fl/fl mice (termed LAT1^-/-^ mice). Two weeks after immunisation with mBSA, tamoxifen was administered to delete LAT1 (Fig S4B–D) and mBSA was injected intra-articularly. There was a highly significant attenuation of arthritis in LAT1^-/-^ mice compared to WT mice (Fig 7H, I). In addition, there was a decrease in the number and proportion of Th1 cells in the knee joints of LAT1^-/-^ mice compared to WT controls, whilst Tregs and Th17 cells were unchanged (Fig 8J).

## DISCUSSION

In this study we showed by immunohistochemistry that LAT1 is expressed in the synovium of RA patients, predominantly by CD4^+^ T cells. This is consistent with publicly available scRNA-seq analysis of human RA synovial biopsies (20) showing abundant *SLC7A5* expression in lymphocytes, with a lower level of expression in monocytes and fibroblasts (Fig. S1). There was found to be a strong correlation between LAT1 expression and serological and clinical markers of disease severity and LAT1 expression aligned closely with a subset of patients in which the inflammatory cell infiltrate is rich in lymphocytes and myeloid cells, as well as NK cells, plasmacytoid dendritic cells, plasma cells and ectopic lymphoid structures (15).

LAT1 deletion has previously been reported to inhibit T cell differentiation and proliferation (11, 21) and the physiological importance of this finding was confirmed here by the fact that its deletion in T cells significantly attenuated arthritis severity, accompanied by reductions in numbers of IFN-γ and TNF-α producing CD4^+^ T cells, whilst numbers of Treg cells remained constant. Genetic deletion of LAT1 in myeloid cells had no effect on arthritis severity, macrophage polarisation or TNF-α and pro-ILβ production *in vitro*, indicating redundancy of LAT1 in these cells.

The altered gene expression profile observed in LAT1^-/-^ CD4^+^ T cells obtained from inflamed joints suggested alterations in TCR/CD28 signalling therefore we examined the capacity of these cells to form functional immune synapses. LAT1 deficient CD4^+^ T cells exhibited impaired actin polarization and reduced recruitment of phospho-tyrosine signalling molecules and CD3ζ to the immune synapse. However, it was particularly striking that LAT1 deficient CD4^+^ T cells from the draining lymph nodes of the same arthritic mice exhibited normal immune synapse formation and TCR signalling. This context specific role for LAT1 is of considerable translational significance as it raises the possibility that LAT1 inhibition may suppress pathogenic T cell activity in the joint, whilst sparing protective T cell activity in lymph nodes. Such considerations are particularly important in the context of chronic inflammatory diseases, where therapy may need to be long-term.

Transcriptional analysis revealed altered expression of genes linked to PI3K/AKT, NFAT and NF-κB signalling pathways in the absence of LAT1, all of which are likely contributors to altered T cell function observed in this study. The PI3K/AKT/mTOR pathway, in particular, contributes to the activation, survival and proliferation of T cells (22, 23) and mTORC1 is known to be sensitive to leucine depletion (24). Indeed, our data highlighted decreased expression of Akt in LAT1 deficient T cells which confirms modulation of this pathway. Another potential contributor to altered T cell activity is the reduced NFATc2 expression observed in LAT1 deficient CD4^+^ T cells (Fig 5C). Consistent with our findings, previous studies in experimental autoimmune encephalomyelitis reported reduced expression of IFN-γ and TNF-α with no inhibition of IL-17A in T cells from NFATc2 deficient mice (25). This implicates impaired NFATc2 signalling as an important mechanism by which LAT1 modulates CD4^+^ T differentiation.

Finally, we studied the effect of LAT1 inhibition in arthritis in the CIA and AIA models. We used a highly selective small molecule inhibitor, JPH203, which has undergone phase I studies for solid tumours (26) and is well tolerated in man. Treatment of mice with already established disease with JPH203 resulted in marked reductions in clinical and histological severity of arthritis, without adverse effects. Similar results were seen with LAT1^-/-^ mice. It was concluded that LAT1 is a druggable target and that this approach is effective in treating ongoing arthritis.

In conclusion, we have established a role for LAT1 in enabling CD4^+^ T cell activation in the inflamed joint and confirm its validity as a therapeutic target in animal models. We propose that LAT1 inhibitors offer a promising therapeutic approach for human disease.

## Acknowledgements

We are grateful to Katja Simon (University of Oxford) for providing Vav-iCre mice, Fiona Powrie (University of Oxford) for providing CD4-Cre mice, Irina Udalova (University of Oxford) for providing LysMCre mice and Tonia Vincent (University of Oxford) for providing UBC-Cre-ERT2 mice.

## Patient and public involvement

Patients and/or the public were not involved in the design, conduct, reporting or dissemination plans of this research.

## Patient consent for publication

Not applicable.

## Ethics approval

Studies involving human participants were reviewed and approved by the West Midlands - Black Country Research Ethics Committee (12/WM/0258) and all subjects provided written, informed consent.

## Data availability statement

The RNA-seq data discussed in this paper have been deposited in NCBI’s Gene Expression Omnibus and are accessible through GEO Series accession number GSE168750 (https://www.ncbi.nlm.nih.gov/geo/query/acc.cgi?acc=GSE168750). All other data are available in the main text or the supplementary materials.

## METHODS

### Mice

All experimental procedures were approved by the Clinical Medicine Animal Welfare and Ethical Review Board and the UK Home Office, in accordance with the 1986 Animals (Scientific Procedures) Act. Mice were housed in individually ventilated cages, maintained at a 21°C ± 2°C and a 12-hour light/12-hour dark cycle, with food and water available ad libitum. LAT1 floxed mice (Slc7a5fl/fl) {Sinclair, 2013 #2924} were purchased from the Jackson Laboratory and bred in-house. Vav-iCre mice were provided by Professor Katja Simon (University of Oxford), CD4-Cre mice were provided by Professor Fiona Powrie (University of Oxford), LysMCre mice were provided by Professor Irina Udalova (University of Oxford) and UBC-Cre-ERT2 mice were provided by Professor Tonia Vincent (University of Oxford). All genetically modified mice were bred on the C57Bl/6 background. Cre mice were crossed with LAT1 floxed mice to delete LAT1 in hematopoietic cells (Vav-iCre), T cells (CD4-Cre) or myeloid cells (LysMCre). Global deletion of LAT1 in adults was achieved by tamoxifen administration of UBC-Cre-ERT2 mice. Two weeks after immunisation, tamoxifen was administered for 5 days by oral gavage (1 mg/day) and sacrificed 8 and 10 days after the last dose for flow-cytometric and histologic analysis, respectively. Successful cre-mediated deletion was confirmed at the end of each experiment by Western blotting, qPCR or using the kynurenine uptake assay. Cre-negative littermates were used as controls. To study the effect of JPH203 on arthritis, C57BL/6 and DBA/1 were purchased from Envigo.

### Antigen-induced arthritis (AIA)

Male or female mice were immunised subcutaneously with 100 μg of methylated bovine serum albumin (mBSA) emulsified with an equal volume of complete Freund’s adjuvant (CFA; BD Biosciences). 21 days later, 100 μg of mBSA was administered by intra-articular injection into the right knee joint while the left knee joint received PBS, as a control. Mice 8-12 weeks old and of a single sex were used in each experiment. Changes in knee swelling were determined by comparing to measurements made prior to intra-articular injection. Where indicated, animals received JPH203 (50 mg/kg per day) or vehicle intraperitoneally from day 1 post arthritis induction for 6 days.

For histological assessment, knees were collected on day 7 and fixed in 10% neutral buffered formalin (Merck) followed by decalcification with 20% Formic Acid for 1 week or 0.5M EDTA for 2 weeks. Decalcified knees were processed into paraffin and embedded coronally. 5 μm thick sections were cut, stained with Safranin-O and Fast green. Histopathologic changes were determined by assessing proliferation of the synovial lining, sub-synovial inflammation, loss of cartilage matrix, chondrocyte death, bone marrow hypercellularity and bone erosion. Each of these 5 parameters was scored by a blinded researcher on a scale of 0-4, giving a maximum possible score of 20. On day 1 post mBSA challenge, mice were given 4 nmol ProSense 750 FAST imaging probe (PerkinElmer) by intravenous injection. 20 hours later, the mice were imaged using the IVIS Spectrum (Perkin Elmer) with an excitation wavelength of 745 nm and an emission wavelength of 800 nm, automatic exposure, and medium binning. Images were analysed using Living Image 4.7 software (Perkin Elmer) to obtain the average fluorescence intensities of a circular region of interest encompassing the knee joint.

Changes in weight distribution were measured using the Dynamic Weight Bearing test (DWB, Bioseb). A single mouse was placed in the device and the weight distribution on each of the four paws of the mice over a duration of 4 min was measured by the high precision force sensors embedded in the floor of the device. Data is presented as a ratio of weight bearing on mBSA injected limb to the PBS injected limb.

### Collagen-induced arthritis (CIA)

Male DBA/1 mice were immunised subcutaneously with 200 μg of bovine type II collagen emulsified in CFA (BD Biosciences) at the base of the tail and on the flank. After immunization, the mice were monitored daily for arthritis. Once an animal showed signs of arthritis, it was randomly assigned to a treatment group and monitored daily. Animals received JPH203 (50 mg/kg per day) or vehicle intraperitoneally and were treated until day 10 of arthritis. Arthritis severity was scored by an experienced, blinded investigator as follows: 0 = normal, 1 = slight swelling and/or erythema, 2 = pronounced swelling, and 3 = ankylosis. All four limbs were scored, giving a maximum possible score of 12 per animal. On day 10, the arthritic paws were collected and fixed in 10% neutral buffered formalin and decalcified with EDTA. Decalcified paws were processed into paraffin and embedded coronally, sectioned and stained with haematoxylin and eosin. Each joint was scored by a researcher blinded to the study as follows: 0 = normal, 1 = cell infiltration with no signs of joint erosion, 2 = inflammation with the presence or erosions limited to discrete foci, and 3 = severe and extensive joint erosion with loss of architecture. Three hind paw joints (proximal interphalangeal, metatarsal phalangeal, and tarsal metatarsal) were scored, and a mean was calculated, giving a maximum possible score of three per animal.

### Isolation of cells from knee joints

The skin was removed from the whole hind limb and most of the muscle was trimmed from the femur and tibia leaving about a millimetre of the muscle immediately surrounding the synovium and the patella. The tibia and femur were dissociated from the surrounding ligament and cartilage without breaking the bone and exposing the bone marrow. The resulting tissue pieces were minced with a pair of scissors and incubated in 500 μl RPMI 1640 (Gibco) supplemented with DNase I (0.1 mg/ml; Sigma) and Liberase TL (0.4 mg/ml; Roche) for 30 min at 37°C in a shaking water bath. Digested soft tissue from the paw joint was then passed through a 70 μm strainer to obtain a single-cell suspension, which was stained for flow cytometry.

### Flow cytometric analysis and cell sorting

For analysis of extracellular markers, cells were stained with Zombie Fixable Viability dye (BioLegend) and unlabelled anti-CD16/32 (BioLegend) to block nonspecific staining in FACS buffer containing PBS with 0.1% BSA and 2 mM EDTA for 15 min in the dark at 4°C. Cells were washed and labelled with fluorochrome conjugated antibodies against cell surface markers in FACS buffer for 30 min in the dark at 4°C. Cells were washed twice and incubated in fixation solution (BD) for 15 min at room temperature. Cells were washed and re-suspended in PBS prior to acquisition. For sorting, extracellular staining of knee cell suspensions was carried out as described above. CD4^+^ T cells were sorted using a FACS ARIA III (BD) with 70-μm nozzle. For immune synapse experiments, CD4^+^ T cells were sorted into 0.5 ml RPMI medium with 5% FCS. For RNA-Seq, cells were sorted into 350 μl RLT buffer with β-mercaptoethanol.

For the kynurenine uptake assay {Sinclair, 2018 #2933}, extracellular staining of cells was performed as described above after which cells were washed with Hank’s balanced salt (HBSS, Sigma) pre-warmed to 37 °C. Cells were then incubated in pre-warmed kynurenine (200 μM in HBSS, Sigma) at 37 °C for 4 min. The reaction was stopped by adding paraformaldehyde (PFA) to a final concentration of 1%. Cells were fixed for 15 min at room temperature, washed twice with FACS buffer and re-suspended in PBS prior to acquisition. For intracellular proteins, cells were stained as above then fixed and permeabilised using the FoxP3/transcription factor staining buffer set (eBioscience) according to the instructions provided. Cells were then washed and stained for intracellular markers in permeabilisation buffer for 45 min in the dark at 4°C. Prior to acquisition, cells were washed twice with permeabilisation and resuspended in PBS. To detect T cell cytokines, cells were stimulated with phorbol 12-myristate 13-acetate (0.02 μg/ml), 0.4 μM ionomycin, and brefeldin A (1.25 μg/ml) for 4 hours prior to staining. For LAT1 detection, cells were incubated with viability dye and FC block followed by fixation and permeabilisation as above. Intracellular staining with anti LAT1 was carried out followed by staining with Alexa Fluor 647 labelled chicken anti-rabbit as a secondary antibody.

Samples were acquired on a Canto II or LSR II or LSR-Fortessa (BD) and analysed using FlowJo Software. All antibodies used for surface and intracellular staining are listed in Table S1.

### Cryosectioning and immunofluorescence staining of mouse knees

Mouse knee joints were obtained immediately after euthanisation and fixed by shaking at 4°C in ice-cold PLP buffer containing phosphate buffer, 4% paraformaldehyde, 1.3% L-lysine and 0.2% sodium periodate. After 24 hours knee joints were washed with PBS and decalcified by incubating in 0.5 M EDTA with shaking at 4°C for 4 days. The samples were then immersed in 30% sucrose in phosphate buffer for another 24 hours. The knee joints were embedded in an 8% gelatin solution supplemented with 20% sucrose and 2% PVP. Samples were then sectioned at 20 μm thickness with a Leica CM3050 cryostat and allowed to air-dry. For immunostaining tissue sections were first air-dried at room temperature for 40 min then at 65°c for 20 min and hydrated with PBS containing 0.3% Triton X-100 for 10 min. Non-specific binding sites were blocked with blocking buffer (10% fetal bovine serum in PBS containing 0.3% Triton X-100) at room temperature after which samples were incubated overnight at 4°C with PE rat anti-CD45 (1:100, clone 30-F11, eBioscience) diluted in blocking buffer. Sections were washed 3 times (5 min each) in PBS solution and the nuclei were counterstained with Sytox blue (Invitrogen) and flat-mounted on microscope glass slides with Fluoromount-G (Invitrogen). Images were acquired on a Zeiss laser scanning microscope 880 using the tile scan function and stitched with a 10% overlap using Zen Black (version 3.1, Zeiss) software. Z-stacks of images were analysed with Imaris software (version 9.5.0, Bitplane).

### Immunohistochemistry and immunofluorescent staining of human synovial tissue

5 μm thick formalin fixed paraffin sections were incubated in a 60°C oven to melt paraffin. Slides were then were deparaffinized in xylene two times for 5 mins each and dehydrated in graded ethanol solutions. After rinsing twice with PBS, sections were incubated in with PBS containing 0.3% Triton X-100 for 10 min and washed with PBS. Citrate and Tris-based antigen-retrieval solutions (pH 6 and 9 respectively) were used for antigen retrieval and non-specific binding was blocked with blocking buffer (3% BSA, 1x PBS, 10% donkey serum; Sigma-Aldrich).

For IHC, the sections were incubated overnight at 4°C with anti-LAT1 (clone 4A2, J-Pharma), washed 3 times with PBS then endogenous peroxidase activity was blocked with 0.3 % H_2_O_2_. Slides were then incubated with biotinylated anti-mouse IgG, for 2 h (Vector) followed by streptavidin-peroxidase complex (Vector) for 30 min then 3,3’-diaminobenzidine (Abcam) for 5 min. The sections were counterstained with hematoxylin and imaged on Leica Aperio ScanScope. LAT1 staining was scored in a blinded manner using the following scoring method: 0 = negative staining, 1 = positive staining of single or limited groups of cells in either the lining or sublining layer, 2 = continuous positive staining of the cells in either the lining or sublining layer and 3 = positive staining of cells in both the lining and sublining layer.

For immunofluorescence staining, slides were stained with DAPI (ThermoFisher) and a background image of the slides was acquired on the GE Cell DIVE system with a 10X objective. Images for the Fluorescein isothiocyanate (FITC), Cyanine-3 (CY3), and Cyanine-5 CY5 channels were also collected to determine the autofluorescence signal in these channels and the autofluorescence from the background image was subtracted from the image acquired after antibody staining. In the first staining round, slides were stained with Anti-Clic5, Anti-LAT1 and Anti-CCR6 for 1 hour at RT followed by 3 washes with PBS for 5 minutes each. Slides were then stained with secondary antibodies in the dark for 1 hour at RT and after washing, the slides were mounted and imaged. Dye inactivation was performed twice by incubating the slides in NaHCO_3_ (0.1M, pH 11.2) and 3% H_2_0_2_ for 15 minutes and DAPI staining and imaging was repeated as above, to create a new background image and assist in image alignment and multiplexing. A second round of staining and imaging was carried out using Anti-CD4, Anti-CD68 and Anti-CD8. Please note that in this manuscript, only Anti-LAT1, Anti-CD4 and Anti-CD68 staining is shown.

### Mouse T cell activation and differentiation

Single cell suspensions were prepared from the lymph node and spleen of age-matched wildtype or knockout mice and CD4^+^ T cells were isolated using the CD4^+^ T Cell Isolation Kit (Miltenyi Biotec). Cells were activated with plate-bound anti-mouse CD3 (5 μg/ml; clone 145-2C11, and soluble antimouse CD28 (2 μg/ml; clone 37.51, eBioscience) for the required duration. Cells were cultured in RPMI medium supplemented with 10% FBS, 50 μM 2-mercaptoethanol and 10,000 U/ml penicillin/streptomycin in the presence of IL-2 (10 ng/ml, PeproTech). For T cell differentiation, the following conditions were used: Th1 (10 ng/ml IL-12, 10 ng/ml IL-2, 5 μg/ml anti-IL-4); Th17 (50 ng/ml IL-6, 10 ng/ ml IL-1β, 10 ng/ml IL-23, 5 ng/ml TGF-β, 5 μg/ml anti-IL-4, 5 μg/ml anti-IFN-γ); Treg (10 ng/ml IL-2 and 5 ng/ml TGF-β). After 4 days of activation, the differentiated T cells were re-stimulated for 4 h with PMA and ionomycin in the presence of Brefeldin A followed by intracellular staining of IFN-γ, IL-17, and Foxp3 to quantify the frequency of Th1, Th17, and Treg cells.

### Mouse bone marrow-derived macrophages

For bone marrow-derived macrophages bone marrow cells were flushed out of tibia and femur from age-matched wildtype or knockout mice. Cells were then cultured in RPMI medium containing 10% FBS and 10,000 U/ml penicillin/streptomycin supplemented with recombinant murine M-CSF (50 ng/ml) for 7 days.

### RNA isolation and quantitative PCR

CD4^+^ T cells were sorted from the arthritic knees as described above. RNA was extracted using the RNeasy Micro Kit (Qiagen), and cDNA was synthesized using the High-Capacity Reverse Transcription kit (Affymetrix, eBioscience). RT-qPCR was performed using TaqMan primers/probes (Slc7a5 Mm00441516_m1, Akt1 Mm01331626_m1; Akt2 Mm02026778_g1; Nfatc2 Mm00477776_m1; Nfkb1 Mm00476361_m1; Nfkb2 Mm00479807_m1, Hprt Mm00446968-m1, Rpl32 Mm02528467_g1, Rpl0 Mm00725448_s1; ThermoFisher Scientific) on a ViiA7 384-well real-time PCR system. All expression levels were normalized to an internal reference gene using the comparative threshold cycle (*C*_T_) method (ΔΔ*C*_T_).

### Quantification of cytokines

Arthritic knees were excised and incubated in 500 μL of RPMI 1640 supplemented with 10% FCS and 10,000 U/ml penicillin/streptomycin at 37 °C in a shaking water bath for 1h. After incubation, the supernatant was collected and centrifuged to remove cells and debris. Enzyme-linked immunosorbent assays (ELISAs) were performed on the clarified supernatants using kits from Invitrogen (TNF-α-88-7324, IL-17-88-7371-88, IL-1β-88-7013-88, GMCSF-88-7334 and IFN-γ-88-8314-88) according to the manufacturer’s instructions. The plates were read using a SPECTROstar Nano microplate reader (BMG LABTECH) at a wavelength of 450 nm. IL-10 and CXCL1 were measured using the Meso Scale Discovery (MSD) platform according to the manufacturer’s instructions.

### Western blotting

Cells were lysed with RIPA buffer (Sigma-Aldrich) supplemented with Halt^™^ Protease and Phosphatase Inhibitor Cocktails (Thermo Scientific) according to the manufacturer’s instructions. Lysates were clarified by centrifugation, and the protein content was determined using Pierce BCA protein assay kit (Thermo Fisher) according to the manufacturer’s recommendation. For immunoblots, equal amounts of each protein sample were separated on a NuPAGE Novex 4-12% Bis-Tris gel followed by conventional blotting and a broad range polypeptide marker (Thermo Fisher) was used to determine the molecular weight of proteins.

### RNA sequencing and bioinformatics

Mouse knees were subjected to enzymatic digestion as described above and CD4^+^ T cells were sorted from the resulting single suspension. Cells were lysed with RLT buffer and RNA was extracted using the Qiagen RNeasy microkit including on-column DNase digestion step, eluted in water and frozen at −80°C. Sequencing libraries were generated using the NEBNext® Single Cell/Low Input RNA Library Prep Kit for Illumina and sequenced on the Illumina NovaSeq6000 platform. Fastq files were mapped to the UCSC Mus musculus genome (mm10) using HISAT2 (v2.1.0; {Kim, 2015 #2615}). Read counts were generated using featureCounts which is part of the Rsubread package (v2.0.1; {Liao, 2019 #2951}) for R (v3.6.3). Statistical analysis of gene counts was carried out using edgeR (v3.28.1, {Robinson, 2010 #2952}) applying TMM normalisation and a 0.05 false-discovery rate p-value adjustment. Bioinformatics analysis was carried out using IPA (Ingenuity Systems, Qiagen, Redwood City, CA, USA). The data discussed in this publication have been deposited in NCBI’s Gene Expression Omnibus {Edgar, 2002 #2953} and are accessible through GEO Series accession number GSE168750: (https://www.ncbi.nlm.nih.gov/geo/query/acc.cgi?acc=GSE168750)

PEAC dataset was analysed using the resource available at https://peac.hpc.qmul.ac.uk/#tab-9442-5

Analysis of RA single-cell RNA-seq data was carried out using the resource available at https://immunogenomics.io/ampra

### Total Internal Reflection Fluorescent Microscopy (TIRFM) imaging

TIRFM imaging was performed with an Olympus IX83 inverted microscope (Keymed) equipped with a 4-line (405 nm, 488 nm, 561 nm, and 640 nm laser) illumination system. The system was fitted with an Olympus 150× 1.45 NA oil-immersion objective and a Photometrics Evolve delta EMCCD camera. Prior to TIRFM imaging, μ-slide 8 well chambered glass coverslips (Ibidi) coated with 0.01% poly-L-lysine (Sigma-Aldrich) were coated with 5 μg/mL of anti-CD3 (clone 145-2C11, Biolegend), 2.5 μg/mL of anti-CD28 (clone 37.51, eBioscience) and 2.5 μg/mL of ICAM-1 (Biolegend) in PBS at 4°C overnight for stimulation of cells. Cells were prepared as indicated earlier and were incubated at 37°C on the μ-slide wells for 15 min, followed by fixation with 4% PFA in PBS for 30 min at RT. After fixation cells were washed three times with PBS, permeabilized and blocked with 0.01% Triton X-100 (Sigma-Aldrich) in blocking buffer (5% BSA/PBS) for 1 h at RT, and then washed three times with PBS. Cell were stained with directly conjugated primary antibodies against anti-CD3ζ-Alexa Fluor-647 (clone 6B10.2, Santa Cruz), anti-pTyr-Alexa Fluor-488 (clone PY20, Biolegend) at 10 μg/mL in 5% BSA/PBS for 1 hour at RT followed by three times washing with PBS. Cells were further stained with CF568 Phalloidin (Biotium) at 1:50 dilution to highlight actin cytoskeleton. Post processing and analysis of the fluorescence images was performed with ImageJ (National Institute of Health). Mean fluorescent intensities of CD3ζ, pTyr and phalloidin were calculated as the sum of intensities in each pixel in the cell spreading area divided by the spreading area of the corresponding cell. The spreading area was determined by thresholding the IRM (Interference Reflection Microscopy) images of each cell.

### Statistical Analysis

Data are presented as the arithmetic mean ± SEM. Comparisons between two groups were carried out using unpaired t tests, Mann-Whitney U tests, one-way ANOVA, or two-way ANOVA depending on the number of groups and nature of comparisons. If significant differences were detected by ANOVA, post hoc Tukey tests (one way) or Sidak’s multiple comparison tests (two way) were completed. Unless otherwise stated, all analysis was completed using GraphPad Prism (GraphPad Software Inc., USA). Probability values (P) of less than 0.05 were considered significant. *P < 0.05, **P < 0.01, ***P < 0.001, and ****P < 0.0001.

**Table S1:** List of antibodies used.

| Antibody | Catalogue No. | Source |
| --- | --- | --- |
| Anti-CD45 | 103137 | Biolegend |
| Anti-CD4 | 12-0041-82 | eBioscience |
| Anti-CD11b | 101217 | Biolegend |
| Anti-Ly6C | 128035 | Biolegend |
| Anti-Ly6G | 127645 | Biolegend |
| Anti-F4/80 | 123147 | Biolegend |
| Anti-CD64 | 139305 | Biolegend |
| Anti-MHCII | 107622 | Biolegend |
| Anti-iNOS | 48-5920-82 | eBioscience |
| Anti-CD206 | 141719 | Biolegend |
| Anti-Arg1 | 12-3697-82 | eBioscience |
| Anti-CD25 | 102035 | Biolegend |
| Anti-Foxp3 | 25-5773-82 | eBioscience |
| Anti-IFN-γ | 48-7311-82 | eBioscience |
| Anti-TNF-α | 554420 | BD |
| Anti-IL-17A | 506928 | Biolegend |
| Anti-Ki67 | 652410 | Biolegend |
| Anti-PD-1 | 135218 | Biolegend |
| Anti-CTLA4 | 106315 | Biolegend |
| Anti-LAG-3 | 125210 | Biolegend |
| Anti-mouse CD16/32 (Fc block) | 101320 | Biolegend |
| Zombie NIR™ Fixable Viability Kit | 77184 | Biolegend |
| Foxp3 / Transcription Factor Staining Buffer Set | 00-5523-00 | eBioscience |
| LAT1 | ab208776 | Abcam |
| Actin | A2228 | Sigma-Aldrich |
| SMC-1 | A300-055A | Bethyl laboratories |
| Anti-LAT1 |  | J-Pharma (clone 4A2) |
| Anti-CD4 | Ab280849 | Abcam |
| Anti-CD68 | NB100-683 | Novus |
| Anti-CD45 | 12-0451-83 | eBioscience |
| Biotin anti-mouse | BA-9200 | Vector laboratories |
| Anti-mouse 647 | A-31571 | Invitrogen |
| Anti-CD3 | 100340 | Biolegend |
| Anti-CD28 | 16-0281-82 | eBioscience |
| Anti-CD28 | 102116 | Biolegend |
| ICAM-1 | 553006 | Biolegend |
| Anti-IL-4 | 504122 | Biolegend |
| Anti-IFN-γ | 505834 | Biolegend |

## SUPPLEMENTARY FIGURES

**Fig S1:**
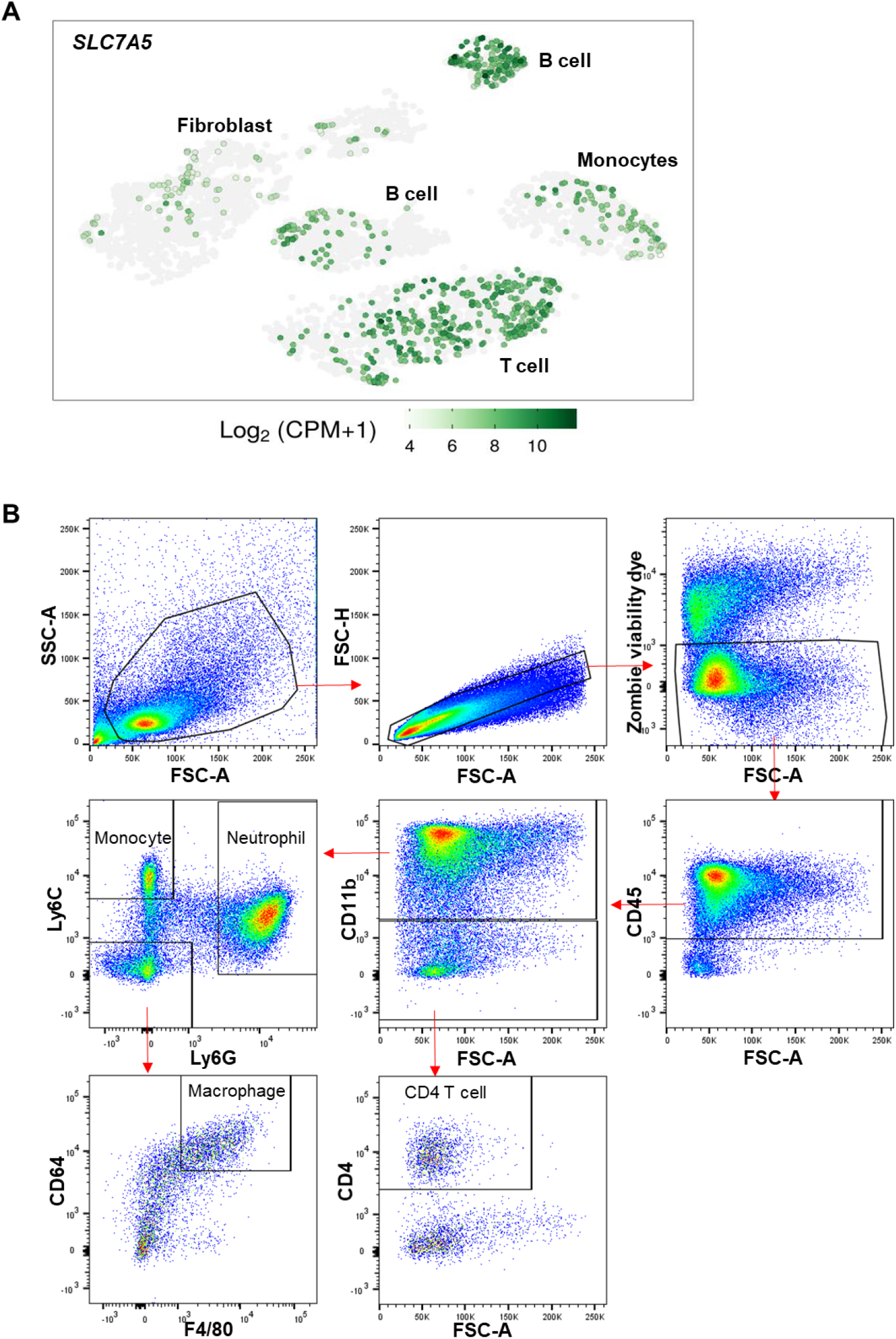
Expression pattern of LAT1 in RA. (A) t Stochastic neighbour Embedding (tSNE) plots of 5265 high-quality cells showing immune cell clusters and LAT1(Slc7a5) expression. Obtained from Zhang et al {Zhang, 2019 #2932} (B) Gating strategy for flow cytometric analysis of immune cells obtained from arthritic mouse joints.

**Fig S2:**
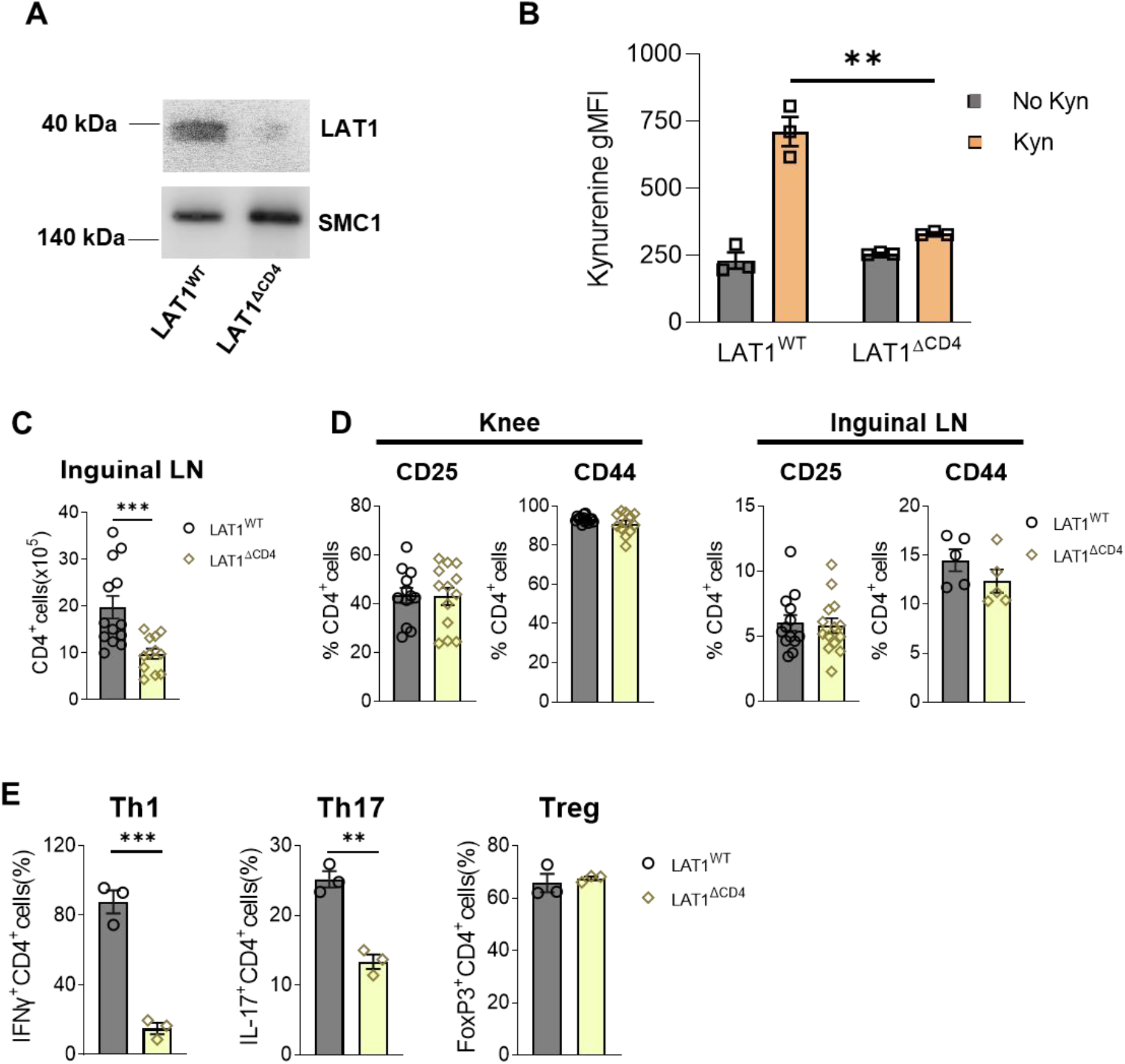
LAT1 deletion in CD4^+^ cells. (A) LAT1 expression detected by western blotting in CD4^+^ T cells isolated from LAT1^WT^ and LAT1^ΔCD4^ mice 48h after TCR stimulation. (B) Kynurenine uptake assay performed on CD4^+^ T cells isolated from LAT1^WT^ and LAT1^ΔCD4^ mice 48h after TCR stimulation. Bars indicate mean ± SD, n = 3 mice from one experiment representative of > 3 separate experiments. (C) Frequency of CD4^+^ T cells in the inguinal lymph nodes of LAT1^WT^ and LAT1^ΔCD4^ mice immunized with mBSA and challenged with mBSA (mean±SEM, n = 12-13 from 3 independent experiments). (D) Frequencies of CD25 and CD44 expressing CD4^+^ T cells in the knees and inguinal lymph nodes of LAT1^WT^ and LAT1^ΔCD4^ mice immunized with mBSA and challenged with mBSA (mean±SEM, n = 12-13 per group, Mann-Whitney U test from 3 independent experiments except for CD44 expression in the LN which depicts data from one experiment). (E) CD4^+^ T cells from LAT1^WT^ and LAT1^ΔCD4^ mice were stimulated under Th1, Th17, or Treg cell skewing conditions and analysed, on day 4, for frequency of IFN-γ^+^ Th1, IL-17^+^ Th17, and Foxp3^+^ Treg cells by flow cytometry. Bars indicate mean ± SD, n = 3 mice from one experiment representative of 3 separate experiments.

**Fig S3:**
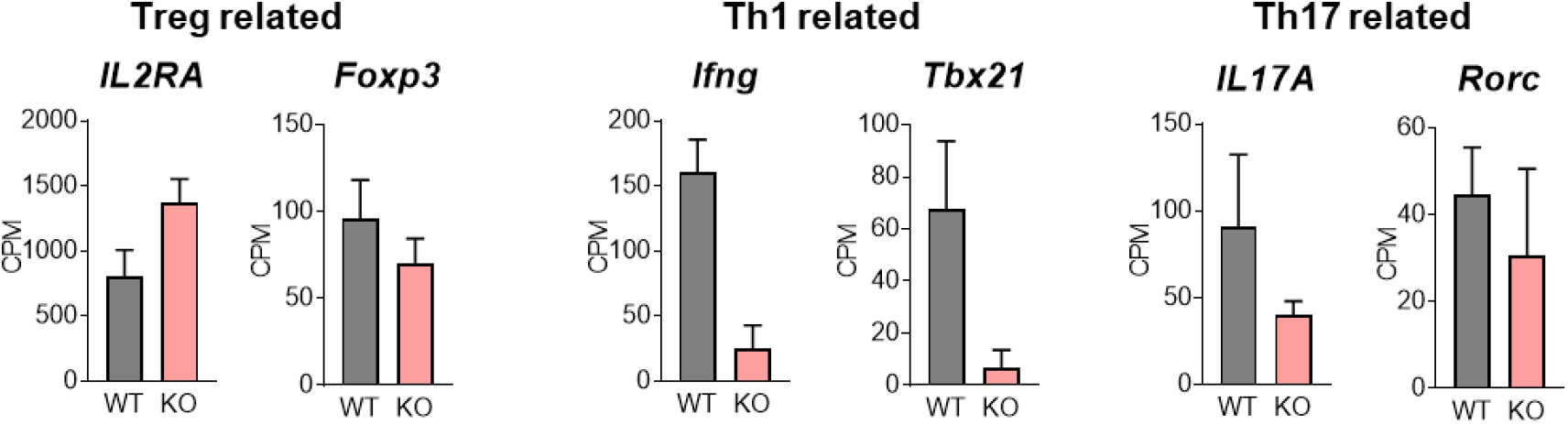
Transcriptome analysis of CD4^+^ T cells from arthritic knee joints of WT and LAT1 KO mice. Bar graphs summarising the mean counts per million (CPM) observed for the indicated genes.

**Fig S4:**
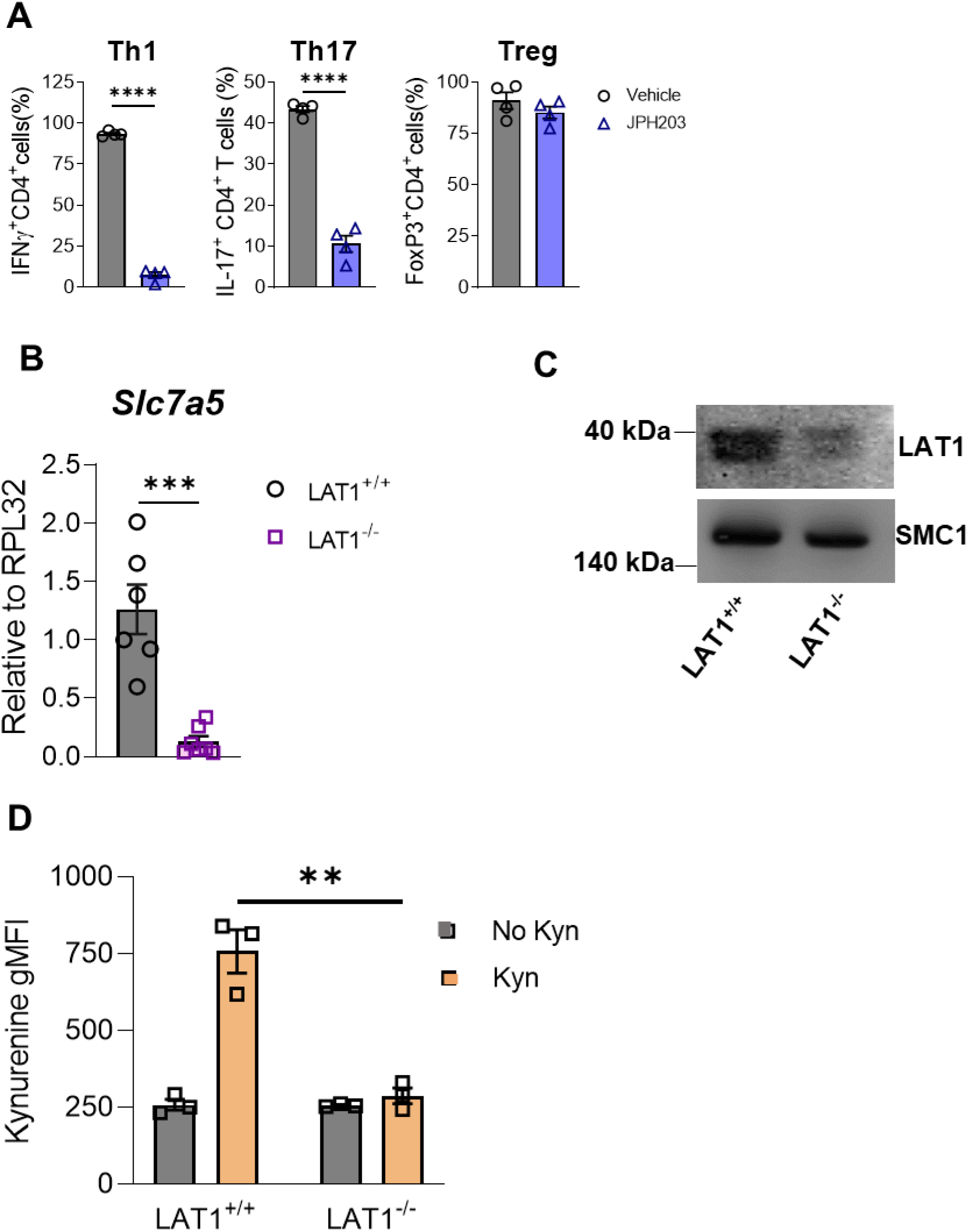
Global deletion of LAT1. (A) CD4^+^ T cells from C57Bl/6 mice were stimulated under Th1, Th17, or Treg cell skewing conditions in the presence or absence of JPH203 (10 μM) and analysed, on day 4, for frequency of IFN-γ^+^ Th1, IL-17^+^ Th17, and Foxp3^+^ Treg cells by flow cytometry. Bars indicate mean ± SD, n = 4 mice from one experiment representative of 3 separate experiments. (B) qPCR analysis of *Slc7a5* mRNA levels in bone marrow cells from LAT1^+/+^ and LAT1^-/-^ mice. Bars indicate mean ± SD, n = 6 mice. (C) LAT1 expression detected by western blotting in CD4^+^ T cells isolated from LAT1^+/+^ and LAT1^-/-^ mice 48h after TCR stimulation. (D) Kynurenine uptake assay performed on CD4^+^ T cells isolated from LAT1^+/+^ and LAT1^-/-^ mice 48h after TCR stimulation. Bars indicate mean ± SD, n = 3 mice from one experiment representative of >3 separate experiments.

